# Cross-catalytic enhancement of peptides and RNA from a common prebiotic activated intermediate

**DOI:** 10.1101/2024.10.02.616211

**Authors:** Raya Roy, Anupam A. Sawant, Sudha Rajamani

## Abstract

Previous origin of life studies have demonstrated the presence of amino acids and nucleotides in the same prebiotic milieu. In this study, we set out to understand the interplay of amino acids with linear or cyclic nucleotides under prebiotically pertinent reaction conditions, especially for its implications for biomolecular evolution. We characterized the cross-catalytic effect of oligomerization, potentially stemming from the simultaneous presence of these two biochemically important monomers. Qualitative and quantitative analysis indicated the formation of longer AMP oligomers and peptides, with 8-10 fold increase in specific reaction scenarios, when compared to reactions that evaluated the monomer oligomerization in isolation. The reason behind such an increase in yield and length, in case of both the oligomers, was the formation of a reactive intermediate. This aminoacylated-AMP (AMP-aa) resulted from a condensation reaction between the nucleotide and the amino acid. We extended this to other amino acids with different R chain characteristics, to comprehend the properties required for the formation of AMP-aa under our reaction conditions. The nonenzymatic formation of these aminoacylated AMP, which in turn resulted in longer oligomers, indicates the plausibility of the emergence of initial steps involved in a primordial translation system.

## Introduction

Peptide bonds and phosphodiester linkages tether amino acids and nucleotides, respectively, resulting in the enzymatic formation of proteins and RNA, thereby playing a vital role in the synthesis of biomolecules in extant biology. Understanding the origin of life primarily relies on characterizing the nonenzymatic formation of these bonds that would have resulted in the build-up of catalytic and informational oligomers in the prebiotic environment; steps central to the transition from chemistry to biology on the early Earth. However, forming these bonds without activated molecules [1-2] or enzymes [3-5] is challenging due to thermodynamic and kinetic barriers [6-8]. Previously studies have shown that subjecting amino acids and nucleotides to high temperature conditions, results in the production of short peptides [9] and RNA [10], respectively. However, one major drawback is that while the formation of peptides is restricted to extreme pH conditions, the abiotic formation of phosphodiester bonds under these conditions results in hydrolysed products. In the context of an RNA peptideWorld Hypothesis [11], preventing hydrolysis and ensuring the accumulation of peptides and RNA becomes imperative while delineating the formation of the first catalytic and informational polymers on the prebiotic Earth.

Recent research in prebiotic chemistry has focused on the role of crowded molecular environments and the interactions between co-solutes to discern the implications of pertinent reactions of interest. Crowding has been shown to influence reaction rates, the putative chemistry that becomes feasible [12-14], leading to the formation of new molecules [15]. These interactions have been shown to aid in molecular survival under prebiotically relevant selection pressures [16] and accelerating the evolution of molecular complexity, with direct implications for the origin of life. AIong similar lines of thought, earlier studies suggest that RNA and peptide precursors could have formed simultaneously [17] under putative prebiotic Earth conditions, with nucleoside phosphates [18-21], cyclic monophosphates [22-23], and amino acids [24-26] coexisting in the same environments. Nonetheless, nucleoside triphosphates (NTPs) have been poorly studied in the aforementioned regard. An important NTP is adenosine 5’ triphosphate (ATP), which is both the primary energy currency in biology and the conventional monomer for RNA synthesis. Its high-energy phosphoanhydride bond facilitates nucleotide polymerization by releasing pyrophosphate (PPi) [2711] and this could potentially be plausible even in the absence of enzymes. Despite its importance, the nonenzymatic oligomerization of ATP under prebiotic conditions remains largely unexplored.

This study explores whether ATP oligomerization, especially in the presence of co-solutes like amino acids in the surrounding environment, could have aided in the formation of oligonucleotides. Further, their simultaneous presence in the immediate environment could have resulted in interesting interactions, with potential implications for affecting their oligomerization reaction rates and yields. In this backdrop, we set out to characterize the formation of oligopeptides and oligonucleotides in a one-pot reaction at neutral pH and elevated temperatures. The compelling outcome that we observed when both the monomers were present in the same reaction mixture, was a resultant increase in both oligopeptides and oligonucleotides, both in terms of the oligomer length and the yield (which increased significantly by 8-10 fold, especially for the former). To delineate the reason behind the aforementioned results, we characterized the different reactions in detail, while being on the lookout for any reactive intermediate that could have resulted in the observed yields and outcome. Relevantly, we detected the formation of an activated/charged amino acid (AMP-aa) in the reaction mixtures. Similar to how activated amino acids serve as key intermediates in the early stages of translation in extant biology, we hypothesized that the AMP-aa produced in our reaction mixture might function as a reactive intermediate, facilitating the oligomerization of both amino acids and nucleotides under our experimental conditions. We also conducted similar studies with 2⍰,3⍰-cyclic adenosine monophosphate (2⍰,3⍰-cAMP) as the nucleotide counterpart. In addition to cyclic monophosphates’ relevance as a prebiotically relevant, ‘intrinsically activated’ monomer for RNA synthesis under geothermal conditions [28-30], these molecules have gotten renewed attention in the context of several prebiotic reactions in the OoL research circles. Pertinently, in reactions involving 2⍰,3⍰-cAMP and glycine (Gly), we also observed high yields of peptide formation and greater stability of the resultant RNA strands.

Additionally, we investigated the formation of AMP-aa intermediates using various amino acids with distinct side chain properties (e.g., acidic, basic, hydrophobic), as well as assessed AMP-aa formation using a regioisomer variant. Most of the amino acids evaluated produced similar outcomes of enhanced cross-catalytic reaction benefits. Our findings offer insights into the properties required for an amino acid to form AMP-aa intermediates. Notably, this study is among the first to show the formation of homo-oligopeptides (up to tetramers) and oligonucleotides (up to pentamers), using nucleotide triphosphate and amino acids in a one-pot reaction. These results highlight how amino acid-nucleotide interactions in a heterogeneous prebiotic soup, would have cross-catalyzed each other’s’ oligomerization, potentially driving an increase molecular complexity under prebiotic conditions.

## Results

### Catalysis of oligopeptide formation in the presence of ATP

Previous studies demonstrated the formation of peptides from non-activated amino acids under wet-dry cycles at elevated temperatures and alkaline conditions (pH 9.5). Analysis of oligopeptide formation in these reactions revealed a pH dependence, with a decrease in peptide formation occurring between pH 4.5 and 8.5. In order to gain a nuanced understanding of how this process could be influenced in the presence of ATP as a co-solute, reactions were carried out at pH 7 wherein previous findings indicated less favourable outcomes for peptide formation. As a first step, the formation of Gly oligomers (peptides) in the presence of ATP was studied to characterize if the overall yields were being altered in the presence of ATP. Towards this, two different concentrations of glycine were evaluated. 25 mM Gly, abbreviated as L for low concentration, or 75 mM Gly, abbreviated as H for high concentration, was subjected to oligomerization in the presence of 25mM of ATP. This was performed at pH 7 without or with MgCl2 (abbreviated M when present) to also understand the effect of divalent cations on these reactions, which were typically subjected to 24 hours wet-dry cycles, at both 90°C and 130°C.

The overall reactions undertaken were as follows (tabulated in Table 1 for ready perusal): ATP + Gly + MgCl2 (25 mM:25 mM:10 mM), abbreviated as AGM 90L or AGM 130L, depending on the temperature used (90°C or 130°C). Higher concentration of Gly-based reactions contained ATP + Gly + MgCl2 in 25 mM:75 mM:10 mM concentration, which is abbreviated as AGM 90H or AGM 130H depending on the temperature of the reaction. Reactions without MgCl2 contained ATP + Gly in 25 mM:25 mM concentration, abbreviated AG 90L or AG 130L, and ATP + Gly in 25 mM:75 mM concentration, abbreviated AG 90H or AG 130H. Control reactions contained 25mM and 75mM glycine, with or without MgCl2 and the following variations of the reactions were performed: Gly+ MgCl2 (25 mM:10 mM), abbreviated GM 90L or GM 130L; Gly + MgCl2 (75 mM:10 mM), abbreviated GM 90H or GM 130H; Gly at 25mM, abbreviated G 90L or G 130L and Gly at 75mM, abbreviated G 90H or G 130H (Table S1). In a parallel iteration of the main reaction, the ATP was replaced with cAMP and these reactions contained 25mM of 2’-3’ cAMP and 75mM of Gly, which were subjected to wet-dry cycles at elevated temperatures of 90°C and 130°C, at neutral pH conditions (Table 1)., abbreviated as cAG 90 or cAG 130, respectively.

**Table 1:** Reactions performed with their concentration and abbreviations: Reaction experiments performed with the starting reactants and reaction parameters. The nucleotides used were either ATP or 2’,3’ cAMP. The concentration of all nucleotides used were kept constant at 25 mM. The different amino acids used with their respective concentrations are mentioned within brackets (). The concentration of MgCl_2_, wherever used was kept constant at 10 mM. The temperatures used are 90°C or 130°C. The abbreviations used corresponds to the specific reactions conducted and has been kept constant throughout.

| Reaction No. | Nucleotide | Amino acid | MgCl<sub>2</sub> | Temperature | Abbreviations |
| --- | --- | --- | --- | --- | --- |
| 1 | ATP | Gly (25 mM) | + | 90 °C | AGM 90L |
| 2 | ATP | Gly (75 mM) | + | 90 °C | AGM 90H |
| 3 | ATP | Gly (25 mM) | + | 130 °C | AGM 130L |
| 4 | ATP | Gly (75 mM) | + | 130 °C | AGM 130H |
| 5 | ATP | Gly (25 mM) | - | 90 °C | AG 90L |
| 6 | ATP | Gly (75 mM) | - | 90 °C | AG 90H |
| 7 | ATP | Gly (25 mM) | - | 130 °C | AG 130L |
| 8 | ATP | Gly (75 mM) | - | 130 °C | AG 130H |
| 9 | 2',3'-cAMP | Gly (75 mM) | - | 90 °C | cAG 90 |
| 10 | 2',3'-cAMP | Gly (75 mM) | - | 130 °C | cAG 130 |
| 11 | ATP | Lys (25 mM) | + | 90 °C | AKM 90 |
| 12 | ATP | Lys (25 mM) | + | 130 °C | AKM 130 |
| 13 | ATP | Lys (25 mM) | - | 90 °C | AK 90 |
| 14 | ATP | Lys (25 mM) | - | 130 °C | AK 130 |
| 15 | ATP | Asp (25 mM) | + | 90 °C | ADM 90 |
| 16 | ATP | Asp (25 mM) | + | 130 °C | ADM 130 |
| 17 | ATP | Trp (25 mM) | + | 90 °C | AWM 90 |
| 18 | ATP | Trp (25 mM) | + | 130 °C | AWM 130 |
| 19 | ATP | β-alanine (75 mM) | + | 90 °C | AAM 90 |
| 20 | ATP | β-alanine (75 mM) | + | 130 °C | AAM 130 |
| 21 | ATP | (S)-3-Amino-4-methoxy-oxobutanoic acid (25 mM) | + | 90 °C | ABM 90 |
| 22 | ATP | (S)-3-Amino-4-methoxy-oxobutanoic acid (25 mM) | + | 130 °C | ABM 130 |

The formation of glycine peptides in the various reaction mixtures was confirmed using tandem MS analysis. The fragmentation masses obtained from purified controls (**Fig. S2(a-e)**) and the theoretically calculated fragment masses from the collision induced fragmentation spectra of glycine oligopeptides that resulted in the reaction, were found to be within 20 ppm error (**Fig. S3(a-k)**). The molecular characterization of the resultant spectra revealed the formation of Gly peptide species, ranging from two residues (diglycine) to a little over ten residues (decaglycine) in ATP-based reactions. The non-ATP based control reactions at 130°C showed only up to the formation of hexamers (hexaglycine). Subsequently, a thorough quantitative analysis was carried out to characterize the varied length of peptides that resulted in the different reaction variations that we tested (as detailed in the beginning of this section). To do this, a LC-MS Q-TOF analysis-based approach was taken to evaluate the different reaction time points. Area under the peak calculation of the different Gly oligomers was used to quantify the yields between ATP-based and non-ATP based reactions.

The aforementioned quantitative analysis demonstrated that the efficiency of Gly oligomerization increased at elevated temperatures (at 130 °C), especially at higher Gly concentrations (**Fig. 1**). Improvement was evident in both the overall yield of the reaction and the generation of longer oligomers, when compared to the reactions performed at 90°C (**Fig. S4 (a-j)**). Also, in almost all the reaction mixtures tested, longer duration of DH-RH (i.e. 24 hours) resulted in longer peptides when compared to the samples collected during earlier time points (e.g. 1 hour, 2 hours, etc,).

**Figure 1:**
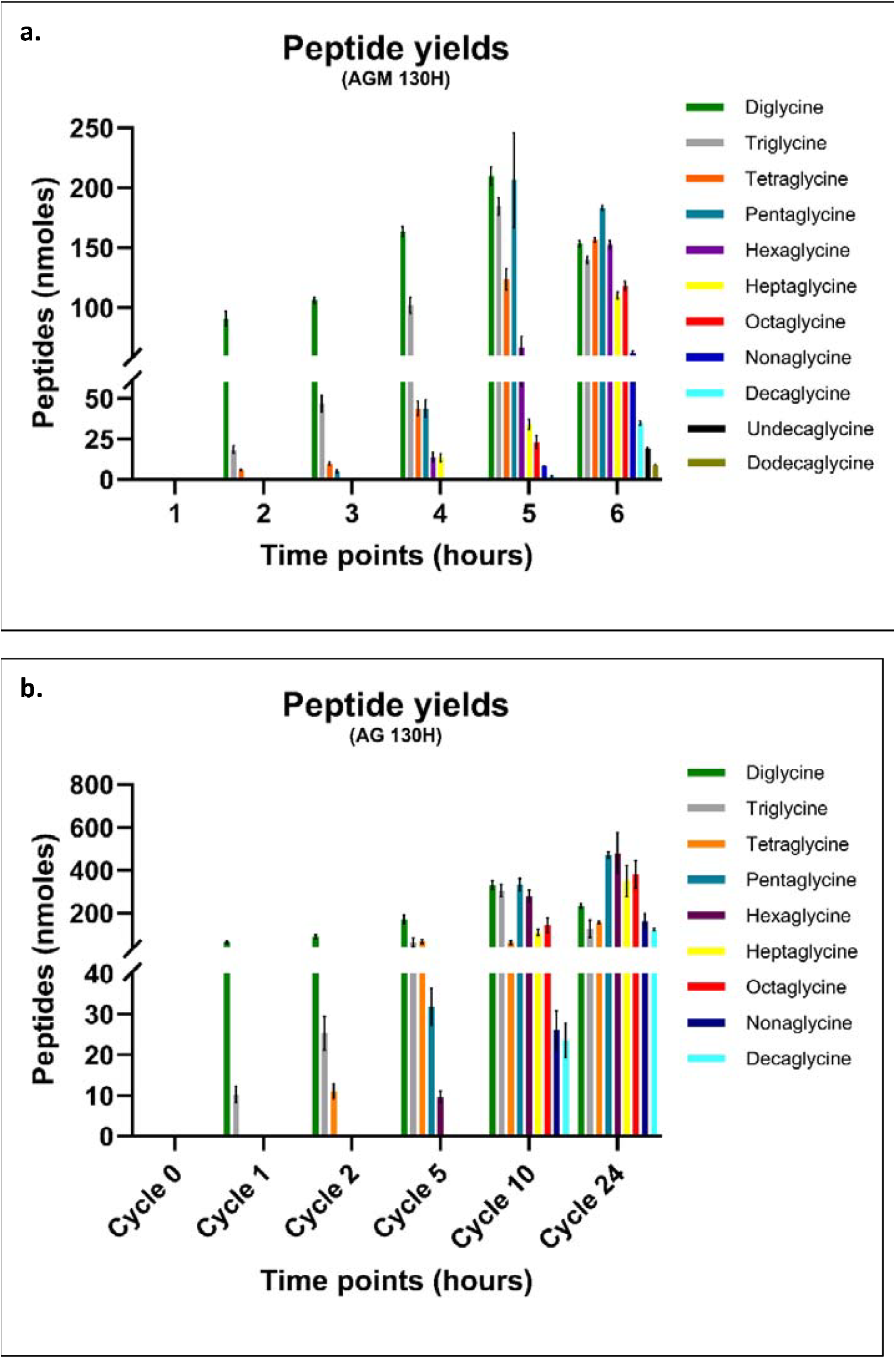
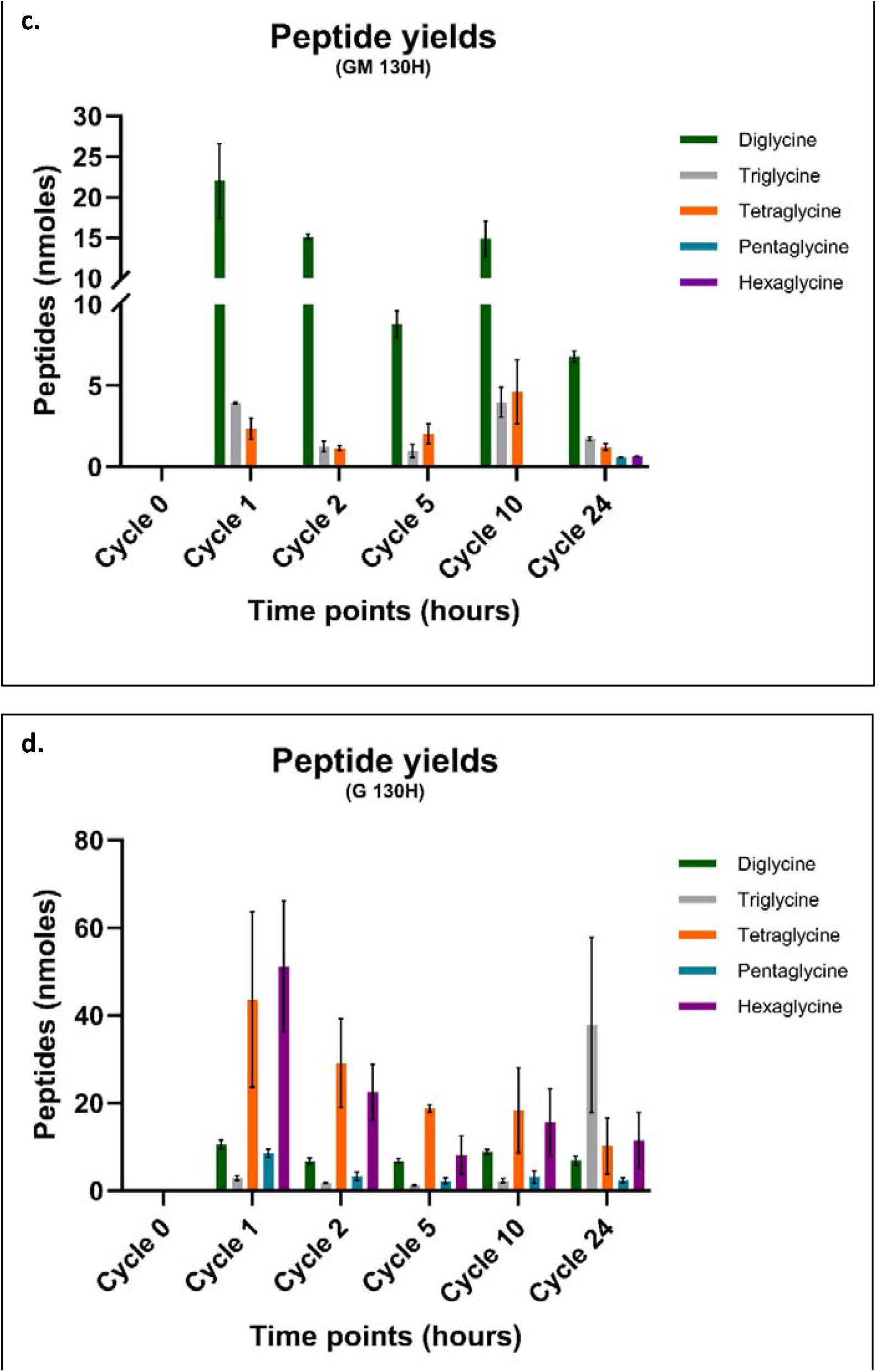
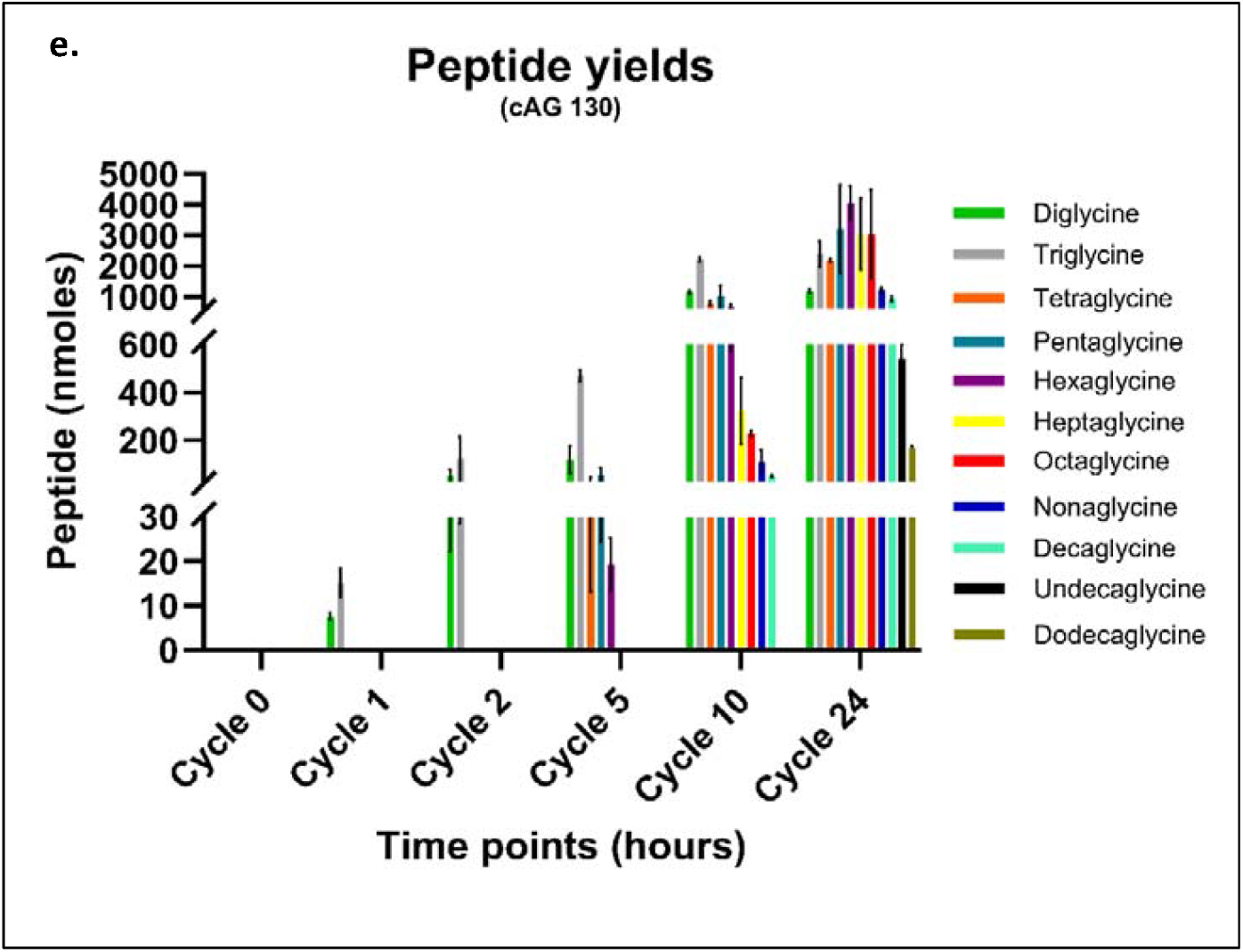
Oligopeptide yields as obtained from mass spectrometry analysis of 130°C. Relative yields of different glycine oligomers formed under wet-dry cycling conditions at 130°C. The Glycine concentration used in the reactions is 75mM (H, high concentration). X-axis indicates the wet-dry cycle time points (0 hr, 1 hr, 2 hr etc., as denoted) at which the samples were analyzed. The Y-axis indicates the relative peptide yields in nmoles (relative to the internal standard used). Reaction conditions: a. ATP + Gly + MgCl2 (AGM 130H), b. ATP + Gly (AG 130H), c. Gly + MgCl2 (GM 130H), d. Gly (G 130H), e. 2’,3’ cAMP + Gly (cAG 130H). The individual bars denote the peptide yields of different length peptides that resulted in the reaction. The error bars represent the standard error of mean for N = 6 replicates.

In the absence of ATP, the 75 mM Gly reactions yielded only up to hexaglycine (Gly)6 (i.e. GM-based and G-based reactions). The cumulative Gly-peptide yields in the 24-hour timepoint of these aforementioned reactions were 10.95 nmoles and 68.7 nmoles, respectively, at 130°C. Relevantly, peptide yields increased by several fold in the presence of ATP, for reactions containing 75 mM Gly. The AGM-based and AG-based reactions yielded up to dodecaglycine (Gly)12 and decaglycine (Gly)10, in 24 hours, yielding cumulative Gly-peptides of 1.11 µmoles and 2.48 µmoles, respectively, at 130°C (24 hours, **Fig. 1(a-d)**). Reactions with 25 mM Gly at 90°C also showed a similar trend, wherein the peptide yield and the resultant length considerably increased in the presence of ATP. In addition, the cAMP-based reactions also resulted in the formation of longer length peptides (dodecaglycine) in the 24th hour cycle, with cumulative Gly-peptide yields of ∼21.3 µmoles (**Fig. 1e**). The most striking observation was on comparison of the net yields; it became evident that the propensity of glycine monomers to result in peptides at physiological pH was clearly enhanced in the presence of ATP as a co-solute. In addition to being a noteworthy result in itself, this also underlines the importance of looking at prebiotic processes from a ‘systems’ perspective to get a realistic understanding of the reaction processes and the product yield distributions in relevant life producing reactions.

### Catalysis of oligonucleotide formation in the presence of glycine

Simultaneous removal of pyrophosphate (PPi) and H2O from ATP molecules results in the formation of oligonucleotides such as nucleotide dimers, trimers etc. The nonenzymatic formation of oligonucleotides could have been affected by co-solutes present in the surrounding milieu as previous studies have shown a prominent affect coming from prebiotically relevant co-solutes like amino acids [15]. Given this, we wanted to ascertain this in the presence of the simplest and, presumptively, the most abundant of amino acid i.e. Gly, to characterize how its presence could have affected oligonucleotides yields. Also, nonenzymatic oligomerization of cyclic mononucleotides has mostly been investigated under alkaline conditions (pH 8-12), with optimal oligomerization occurring at pH 10 [10, 28]. Given this, we wanted to determine what effect the presence of amino acids such as Gly, would have had on oligonucleotide formation. In this backdrop, oligonucleotide formation and yields for both ATP and 2’,3’-cAMP monomers, were characterized at pH 7.

To elucidate the oligomerization efficiency of nucleotides (ATP) in the presence of amino acids, the same reaction setup of AGM and AG reactions were used with varied concentration of Gly (25 mM or 75 mM) **(Table 1)**. In addition to the AGM- and AG-based reactions, control reactions were carried involving 25mM ATP with MgCl2 (abbreviated AM90 or AM130) and 25 mM ATP alone (abbreviated A90 or A130, Table S1). For cAMP-based reactions, a mixture of 25mM 2⍰, 3⍰-cAMP with 75mM Glycine (pH 7) was subjected to wet-dry cycling at 90°C and 130°C (abbreviated cAG90 and cAG130, respectively) **(Table 1)**. The control reaction contained just 25 mM 2’,3’-cAMP (abbreviated cAMP), which was then subjected to similar conditions as the main reaction setup **(Table S1)**.

Anion exchange chromatography was employed to investigate the formation of nucleotide oligomers across the various reactions and at different reaction cycles **(Fig. S5-S8)**. Notably, following ATP elution at around 6.9 min retention time (RT), new peaks were seen to elute at and after RT 7.4 min, indicating potential oligomerization of AMP. Quantitative analysis of these peaks over different time intervals revealed dynamic changes when compared to what was observed in the 0-hour cycle **(Fig. S9)**. Pertinently, the stability of these new peaks appeared to be influenced by the presence of Gly in the reaction mix in addition to the reaction temperature. However, it is essential to acknowledge the limitation of HPLC chromatography in definitively confirming longer nucleotide formation, especially because of its sensitivity to detect oligomers being impacted when they are present at really low concentrations. Interestingly, in AM reactions at 90°C, greater stability was observed in the absence of Gly. Conversely, at 130°C, reactions containing Gly exhibited enhanced product stability.

To understand the aforesaid observations, we conducted a qualitative LC-MS analysis of the various reaction mixtures. The collision-induced fragmentation patterns obtained via TOF-MS/MS indeed confirmed the presence of oligonucleotides in both the Gly-containing and Gly-free reactions **(Fig. S11, S12)**. These were consistent with the fragmentation pattern and the theoretical calculated values that was obtained for purified oligonucleotides (Fig. S10). However, the spectra revealed variations in the lengths of the resultant oligonucleotides. Notably, reactions with Gly (AGM and AG reactions) demonstrated the presence of oligonucleotides of up to four residues (e.g. AMP tetramer), whereas the Gly-free reactions (AM and A reactions) consistently showed oligonucleotides of up to only two residues (e.g. AMP dimer) across different cycles.

Further, the yields of the different nucleotide oligomers formed in the reactions were calculated using the area under the peak analysis feature on LC-QTOF **(Fig. 2)**. Oligonucleotide yields were higher at 130°C than at 90°C **(Fig S14)**, potentially due to the higher thermal energy available at 130°C. The AMP dimer yields were higher in the AM- and A-based reactions than in the AGM and AG reactions and this could be attributed to the AMP dimers potentially being consumed in the formation of longer oligonucleotides in the latter two reactions. Another possible reason for this could be that the ATP molecules were being taken up by Gly to form other reaction intermediates, thereby decreasing the availability of ATP molecules for oligonucleotide formation. This is clearly evident from the detailed mass characterization that we undertook (detailed in the next section).

**Figure 2:**
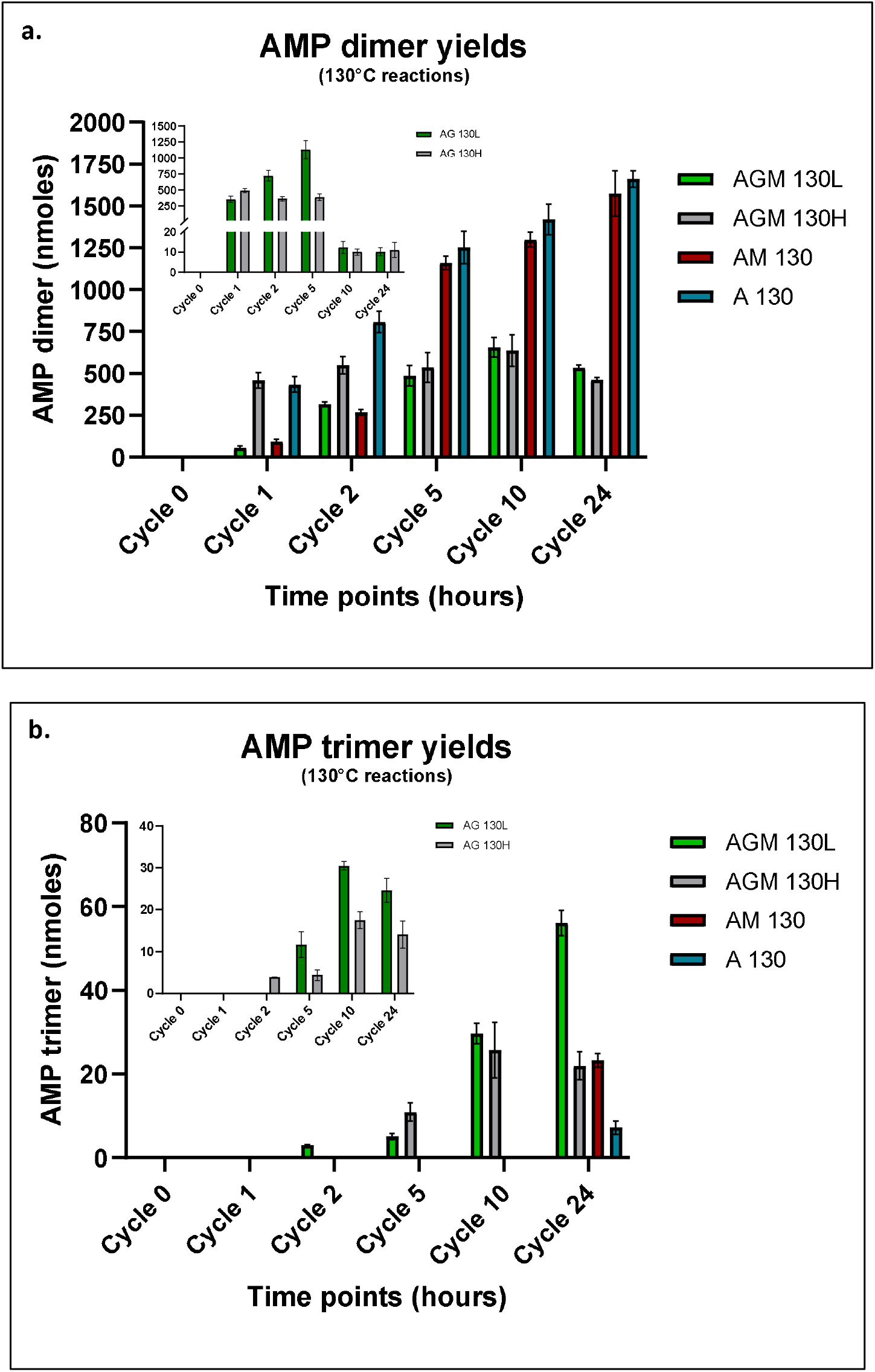
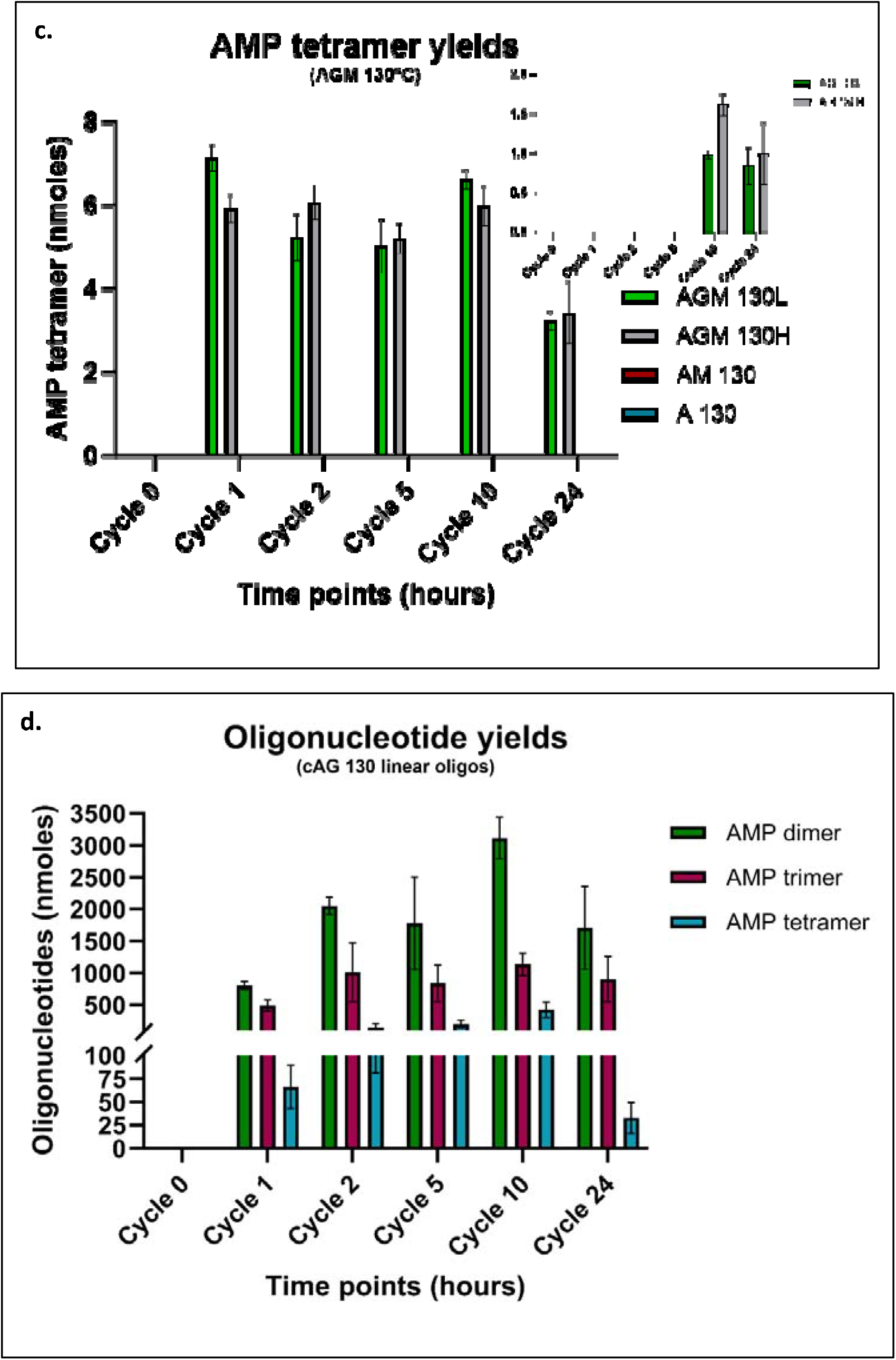
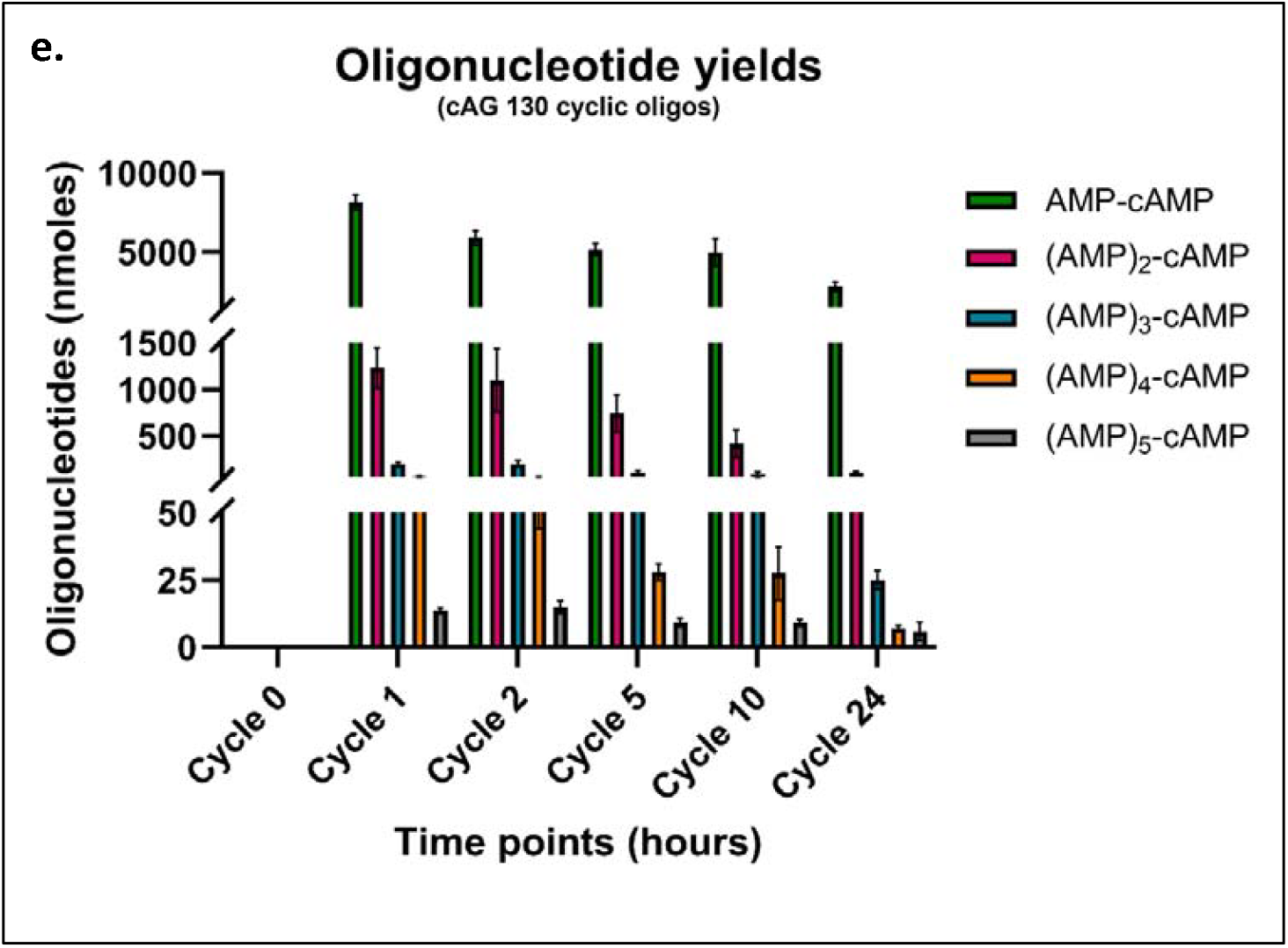
Oligonucleotide yields as obtained from mass spectrometry analysis of 130°C. Relative yields of **a**. AMP dimers, **b**. AMP trimers and **c**. AMP tetramers formed under wet-dry cycling conditions at 130°C and different concentrations of glycine (25 mM, abbreviated as L or 75 mM, abbreviated as H) through different cycles in the various reactions studied. Oligonucleotide yields in reactions involving ATP + Gly+ Mg^2+^ (AGM), ATP+ Mg^2+^ (AM) and ATP (A) mixtures are demonstrated. The insets demonstrate the reaction yields under similar reaction conditions in ATP + Gly (AG) reaction mixtures. Relative yields of different length **d**. linear and **e**. cyclic oligonucleotides formed under wet-dry cycles at 130°C and 75mM of glycine through different cycles in 2’,3’ cAMP + Gly (cAG) reaction mixtures. X-axis indicates the wet-dry cycle timepoints at which samples were analyzed. The Y-axis indicates the relative oligonucleotide yields in nmoles (relative to the internal standard used) of different length oligonucleotide compounds formed. The error bars represent the standard error of mean for N = 6 replicates.

Also, in the cAMP-based reactions, longer oligonucleotide yields were only observed at 130°C and in the presence of Gly. Oligonucleotide formation was not observed in the absence of Gly or at 90°C as was evident from the HPLC chromatograms **(Fig. S13)**. Therefore, only the oligonucleotides from the cAMP-based reactions that were carried out at 130°C were systematically characterized. Two types of oligonucleotides were identified in the cAMP-based reactions: linear oligonucleotides (where all the molecules were linear) and cyclic oligonucleotides (where the last nucleotide remained in the cyclic form). The fragmentation spectra of linear oligonucleotides were similar to those obtained from ATP-based reactions. The theoretically calculated fragment masses and the fragmentation masses obtained from the collision induced fragmentation spectra of cyclic oligonucleotides in the reaction, were found to be within 20 ppm error **(Fig. S15 a-e)**. The longest observed oligonucleotides were the AMP tetramer (AMP-AMP-AMP-AMP) for the linear form and the AMP hexamer (AMP-AMP-AMP-AMP-AMP-cAMP) for the cyclic form. Significantly, the oligomer yields for both the linear and cyclic oligonucleotides quantified from the 2’,3’-cAMP reactions were clearly higher than what was seen in the ATP-based reactions **(Fig. 2g, h)**. The reason could be attributed to the higher stability of cAMP (the starting reactant) at physiological pH than ATP. ATP was more prone to hydrolysis and this was observed in our HPLC time course chromatograms too **(Fig S8 and S13)**.

### Observed intermediate for the cross-catalytic enhancement of oligopeptides and oligonucleotides

To comprehend the elevated yields of oligopeptides in our reactions, and the increased residue length observed in case of the resultant oligopeptides and oligonucleotides, we investigated further to understand if a reactive intermediate may have enabled these outcomes. Using mass analysis, the formation of activated amino acid i.e. AMP-aa was readily detected in our reactions where both the ATP and amino acid were present in the starting reaction mix. The formation of this AMP-aa could have been facilitated by a nucleophilic interaction involving the carboxylic or the amino group of the amino acid and the α-phosphate group of ATP. In all the reactions containing both ATP (25mM) and Gly (25mM or75mM), and in the presence/absence of MgCl_2_ [AGM & AG reactions **(Table 1)**], the formation of AMP-Gly was observed by mass analysis. This indicated that the nucleophilic interaction is feasible regardless of the presence of divalent cations like Mg^2+^.

For qualitative confirmation and molecular characterization, tandem MS was performed and collision-induced fragments were compared with that of the theoretically calculated fragmentation masses obtained for AMP-Gly. Four linkages of AMP-Gly **(Fig. S16)** with similar precursor mass [404.0845 Da] were deduced from the TOF MS/MS fragmentation spectrum **(Fig. S17a)**. The two possibilities for the aforementioned observations could be as follows: the lack of regiochemical control in wet-dry reactions and/or the presence of H atom as the R group of glycine potentially makes it compliant to result in different kinds of linkages. Even though the similar precursor masses meant that they sorted similarly before fragmentation in Q1 chamber, different masses were observed during collision-induced fragmentation. Pertinently, quantitative analysis of our data sets for AMP-aa based peaks ascertained that they occurred in higher yields at 130°C and in the presence of MgCl_2_ than when compared to 90°C and in the absence of MgCl_2_, respectively **(Fig. 3a & S18a)**.

**Figure 3:**
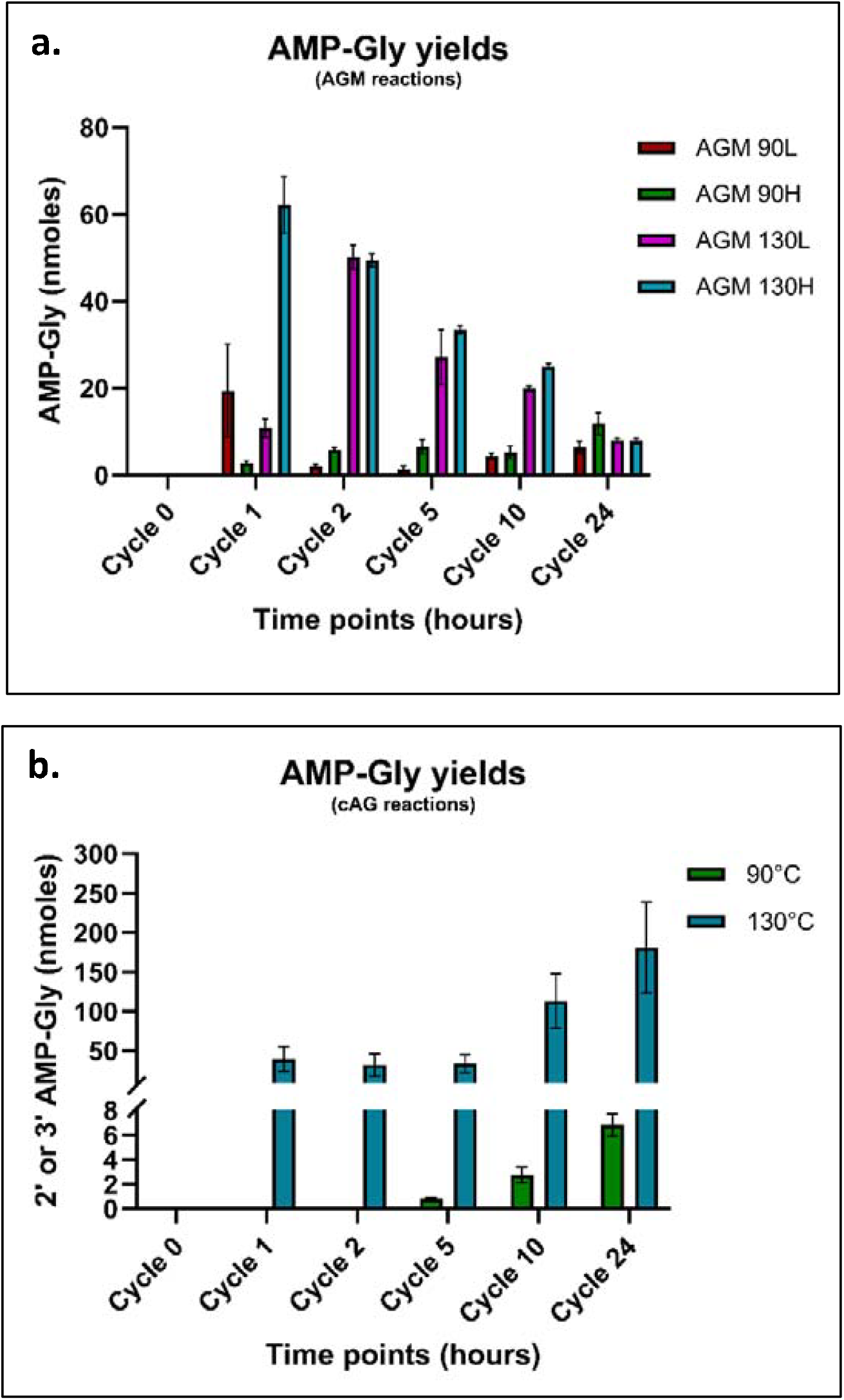
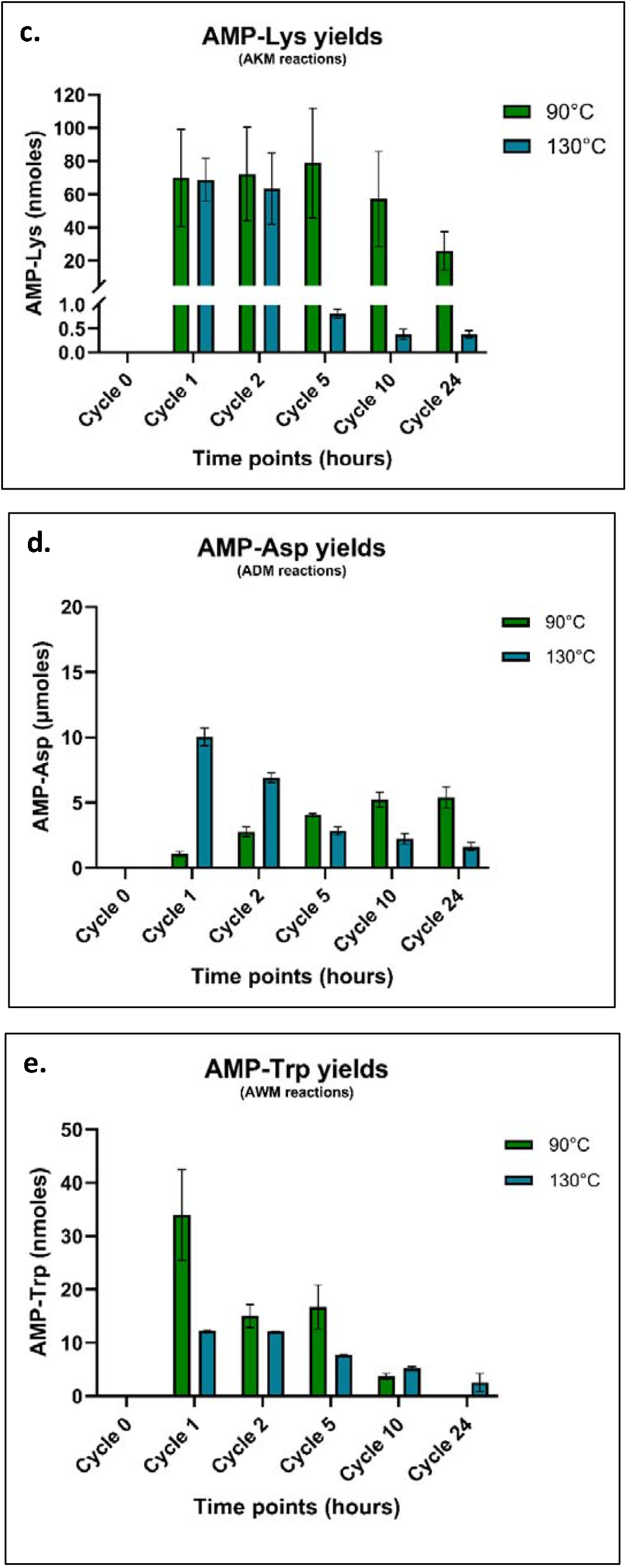
AMP-aa yields as obtained from mass spectrometry analysis. Relative yields of AMP-aa found under wet-dry cycling conditions at elevated temperatures of 90°C & 130°C for various α-amino acids tested. Fig **3a** demonstrates the relative yields (relative to the internal standard) of AMP-Gly found in ATP + Gly+ Mg^2+^ (AGM) reaction mixtures containing two different concentrations of glycine (25 mM, abbreviated as L or 75 mM, abbreviated as H). Similarly, Fig **3b** demonstrates the relative yields of 2’ or 3’ AMP-Gly found in 2’,3’ cAMP + Gly (75mM), cAG reaction mixtures at two different temperature conditions. Fig **3c, d**, and **e** demonstrates the relative AMP-Lys, AMP-Asp, and AMP-Trp found under wet-dry cycling conditions at 90°C & 130°C through different time points in ATP (25mM) + Lys (25mM) + Mg^2+^ (10mM); ATP (25mM) + Asp (25mM) + Mg^2+^ (10mM); ATP (25mM) + Trp (25mM) + Mg^2+^ (10mM) reaction mixtures respectively. X-axis indicates the wet-dry cycle timepoints at which samples were analyzed. The Y-axis represents the relative yields of different AMP-aa compounds found. The error bars represent the standard error of mean for N = 6 replicates.

Further, to explore the implications arising from the differential positioning of the -COOH group relative to the -NH_2_ group in certain amino acids, we used β-alanine to check for the resultant formation of AMP-aa. A mixture of 75 mM β-alanine, 25 mM ATP, and 10 mM MgCl_2_ was subjected to wet-dry cycling at both 90°C and 130°C for 24 hours (abbreviated as AAM 90 and AAM 130, respectively, Table 1). However, qualitative analysis did not reveal the formation of AMP-β -Ala. Furthermore, whether AMP-aa could result with amino acids containing amino groups not only in the α-NH_2_ position but also in the R group was also investigated. For this, 25 mM lysine was mixed with 25 mM ATP, with and without 10 mM MgCl_2_, and was then subjected to wet-dry cycling at temperatures of 90°C and 130°C. These reactions were abbreviated as AKM 90 or AKM 130 for ATP + Lys + MgCl_2_ (25 mM: 25 mM: 10 mM) and AK 90 or AK 130 for ATP + Lys (25 mM: 25 mM), respectively (Table 1). Qualitative analysis by tandem MS indicated the formation of AMP-Lys **(Fig. S17b)**, particularly at a higher yield in the presence of MgCl_2_ **(Fig. 3c & 18b)**. As a next step, we tried to introduce more variation in the side chain, and for this we also evaluated the formation of AMP-aa using aspartic acid (which has a negatively charged R group) and tryptophan (which has a bulky-hydrophobic R group). Asp and Trp, at a concentration of 25 mM each, were subjected to wet-dry cycling along with 25 mM ATP and 10 mM MgCl_2_ at both 90°C and 130°C Reactions were abbreviated ADM 90 or ADM 130 for ATP + Asp + MgCl_2_ at 25 mM: 25 mM: 10 mM; and AWM 90 or AWM 130 for ATP + Trp + MgCl_2_ at 25 mM: 25 mM: 10 mM, respectively (Table 1). Fragmentation spectra obtained from the tandem MS profiles suggested the formation of AMP-Asp **(Fig. S17c)** and AMP-Trp **(Fig. S17d)**, respectively, in the above reactions.

We also determined the quantitative yield that was obtained for the different AMP-aa compounds using a similar approach as mentioned above. At 130°C, all the amino acids exhibited the highest yields of AMP-aa in their respective reaction. However, the relative yields of AMP-aa varied notably among the different amino acids and the different wet-dry cycle time points analyzed. For example, in reactions at 130°C, the highest yield was observed for Asp (6.9 µmoles at cycle 2), followed by Lys (68.6 nmoles at cycle 2), Gly (62.2 nmoles at cycle 1), and Trp (12.3 nmoles at cycle 1) **(Fig. 3 c-e)** This yield difference could be due to the different pKa values of the α-COOH groups in the amino acids. It could also be due to the presence of two -COOH groups in Asp and two -NH2 and one -COOH groups in Lys. In order to delineate this, we used the α-COOH methyl ester version of aspartate ((S)-3-Amino-4-methoxy-oxobutanoic acid). 25 mM of (S)-3-Amino-4-methoxy-oxobutanoic acid was mixed with 25 mM ATP and 10 mM MgCl_2_ and subjected to wet-dry cycles at elevated temperatures of 90°C and 130°C for 24 hours (abbreviated ABA 90 and ABA 130) (Table 1). Qualitative analysis revealed no formation of AMP-Asp when the α-COOH group was protected with a methyl group in Asp as this would have led to reduced nucleophilicity of the amino group, resulting in no formation of AMP-aa. Relevantly, this result could also imply that the α-COOH is the major group participating in the formation of AMP-aa.

Moreover, we were also interested in characterizing if the formation of AMP-Gly was possible when substituting the conventional nucleotide with a cyclic nucleotide. For this, 25 mM 2⍰, 3⍰-cAMP with 75 mM glycine (pH 7) was subjected to wet-dry cycling at 90°C and 130°C, abbreviated cAG 90 or cAG 130, respectively. MS/MS fragmentation spectra confirmed the formation of 2’-AMP-Gly or 3’-AMP-Gly in the cAMP-based reactions **(Fig. S17e)**. Unlike the ATP-based reactions, the formation and accumulation of 2’-AMP-G or 3’-AMP-G was gradual, with the highest yield observed at cycle 24 (∼180 nmoles) **(Fig. 3b)**.

Finally, after confirming the formation of activated amino acids of Lys, Asp and Trp (all naturally occurring α-amino acids), we wanted to check for the formation of oligonucleotides and oligopeptides in these reaction mixtures as well. Oligonucleotides up to tetramers were observed in all the three reactions **(Fig. S19)**, similar to what we saw in glycine-based reactions. Oligopeptides of length trimer, tetramer and pentamer were observed for Trp-, Lys- and Asp-based reactions, respectively, which were confirmed using tandem MS characterization **(Fig. S20)**. Pertinently, precursor masses of complex compounds such as AMP-AMP-G/K/D/W and AMP-AMP-AMP-G/K/D/W were also observed in the AGM, AKM, ADM and AWM reactions, respectively. Just as in AMP-aa, these molecules also contain the high energy mixed anhydride bond between the last nucleotide and the carboxylate group of amino acid. Interestingly, these molecules were found to be present in very low amounts in the reaction mixtures, which could mean that these unstable molecules were being quickly consumed for the formation of AMP trimers and AMP tetramers **(Fig. S21 a-h)**.

## Discussion

This study investigated interactions between amino acids and nucleotides under prebiotically relevant conditions, focusing on characterizing product formation and their yields. On simulating the conditions of terrestrial geothermal environments by employing wet-dry cycles at physiological pH, longer oligomers were produced in higher yields when both the bio monomers were present, when compared to reactions that involved only the oligomerization of the individual biomolecule. The study also observed the formation of a reactive intermediate, ‘activated amino acid’ (AMP-aa), that was likely driving the synthesis of longer oligonucleotides and peptides under our reaction conditions.

In reactions involving ATP and glycine, formation of dodecaglycine and decaglycine was observed, with cumulative peptide yields (of various length peptides) of 1.11 µmoles and 2.48 µmoles resulting in the presence and absence of Mg^2+^, respectively. In contrast, under the same reaction conditions, absence of ATP yielded only up to hexaglycine, with cumulative peptide yields of ∼10.95 nmoles and 68.66 nmoles, in the presence and absence of Mg^2+^, respectively. This clearly demonstrates a pronounced enhancement of both peptide yield and oligomer length, when ATP is included in the starting reaction mixture. Similarly, when Gly was replaced with Lys/Trp/Asp, peptide formation was only observed in the presence of ATP at the pH conditions evaluated. As mentioned earlier, peptide formation using any amino acids has been shown to be feasible only under extreme pH conditions in previous studies. And, this was even more difficult for amino with bulky R groups such as Lys/Trp/Asp. Our study possibly is one of the first ones to report appreciable yields for peptide for amino acids with cationic, anionic and hydrophobic R chains.

We also observed that not all amino acids tested resulted in the formation of activated amino acids. For instance, β-alanine did not form an activated reaction intermediate with ATP This could either be due to the altered nucleophilic properties of the -COOH and -NH_2_ groups when they are separated by a methyl group, or due to the high pKa (3.6) of the - COOH group in β-alanine. Nonetheless, lysine, which has an additional -NH_2_ group, did result in the formation of AMP-Lys **(Fig S17)**. The fragmentation pattern of AMP-Lys also suggested that the resultant bond is from the reaction between the phosphate group of the nucleotide and the α-COOH of lysine. In all, the yields of AMP-aa in the reactions involving the amino acids Gly, Lys, Asp, and Trp are as follows: AMP-Asp >> AMP-Lys > AMP-Gly > AMP-Trp. It is important to note that the pKa values of the α-COOH groups of the amino acids used are: Asp (1.88) << Lys (2.18) < Gly (2.34) < Trp (2.83). This suggests that the formation of AMP-aa could be pKa-dependent.

Since all reactions in this study were conducted at physiological pH, a lower pKa of the α-COOH meant greater ease of activation of amino acids by ATP [32]. Importantly, protecting the α-COOH group of Asp did not yield AMP-Asp in a test reaction that we carried out, thereby confirming the importance of the α-COOH group of the amino acid in the formation of activated amino acids under our reaction conditions. Further, it is important to note that AMP-Gly is formed due to four different kinds of linkages as mentioned in the results section and as has also been reported previously [33]. Amongst these, the phosphoramidate (P-N) bond and the anhydride bond (P-O-C) was found in low yields in our reaction suggesting that they are probably readily consumed due to their higher reactivity [34-36]. On the other hand, the AMP-aa with the ester bond (CO-OR) and the amide bond (CO-NH) were observed in higher yields, resulting in a clean TOF-MS/MS spectra.

As for the role of divalent cations like Mg^2+^ in the formation of peptides, it is somewhat complex. Similar to previous studies, we also observed that the peptide yields in reactions with and without ATP were lower in the presence of Mg^2+^ than when it was absent. This could be because of the Mg^2+^ inhibiting peptide synthesis by chelating with amino acids and catalysing hydrolysis like what has been reported [31]. Alternately, the stability or the accumulation of AMP-aa intermediates, and thereby their yield, is possibly higher in the presence of Mg^2+^. The quantitative yields from mass spectrometry suggested that the AMP-aa intermediates were possibly being consumed more readily in the absence of Mg^2+^ to result in peptides.

In the case of oligonucleotide formation under the reaction conditions studied, those reactions where there were no amino acids resulted in the formation of AMP dimers at much higher yields than what was observed in the presence of amino acids. Importantly, in oligonucleotide oligomerization reactions where amino acids were present, complex intermediates such as AMP-AMP-G/K/D/W and AMP-AMP-AMP-G/K/D/W, as well as oligonucleotides up to tetramers, were observed using mass spectrometry. The phosphoramidate (P-N) bond or the mixed anhydride bond that can form between AMP and amino acids, like in AMP-aa, AMP-AMP-aa, and AMP-AMP-AMP-aa compounds, is quite unstable making it highly reactive [34-36]. During the rehydration phase of the wet-dry cycles, these molecules can interact with AMP monomers or oligomers, to form upto tetramers (AMP)_4_ and hexamers ((MP)_5_-cAMP in ATP and cAMP-based reactions respectively Unlike peptide formation, the oligonucleotide formation was higher in the presence of Mg^2+^ than when it was absent. Similar to its role in extant biological reaction, Mg^2+^ seems to be essential for the formation and stability of RNA even under nonenzymatic conditions [35,36]. In aqueous conditions, Mg^2+^ forms stable complexes with ATP, thereby neutralizing its charge while and also coordinates with and polarizes the pyrophosphate (PPi) within ATP [39-40]. This characteristic of Mg^2+^ likely leads to the formation of the complex compounds that have been mentioned above, while also facilitating the formation of longer RNA strands.

Similar to ATP-based reactions, cAMP-based reactions also showed a positive enhancement in the formation of both peptides and oligonucleotides. The reaction did not require MgCl_2_ since cAMP is an intrinsically active molecule. Additionally, cAMP is highly stable at neutral pH and under the temperature conditions used in our reactions **(Fig S13)**. The ring opening of cAMP, and therefore the formation of peptides and oligonucleotides, was only observed in the presence of amino acids. Reactive intermediates such as AMP-G, AMP-AMP-G, and AMP-AMP-AMP-G were also detected in the cAMP-based reactions. Furthermore, due to the higher stability of cAMP, the yields of both oligonucleotides and peptides were 3 and 6 fold higher than what was observed in the ATP-based reactions.

This study aims to characterize and quantify the products formed from common reactive intermediates, specifically AMP-aa, which emerges from interactions between two prebiotically pertinent co-solutes, i.e. nucleotides and amino acids. Aminoacylation, a key step in translation, is critical to understanding the nonenzymatic formation and utilization of these reactive intermediates. Relevantly, understanding the corresponding prebiotic process also sheds light on the early development of the translation system in modern biology. Recent research on prebiotic aminoacylation and translation has highlighted the significance of tRNA aminoacylation and subsequent peptide formation and amino acid transfer [11,41-44]. However, it is equally important to explore the nonenzymatic counterpart, i.e. single-nucleotide-driven peptide formation, which would have been a hallmark of the RNA-peptide world. Before the evolution of long tRNAs, these single-molecule reactions involving reactive intermediates from essential monomers would have provided a foundation for increasing the complexity of the system. Even today, amino acid activation precedes tRNA activation in cells, suggesting that during the early stages of life, the activation of amino acids to form longer peptides and nucleotides was likely a crucial step toward the origins of the minimal translation machinery.

### Materials

All the amino acids (glycine, tryptophan, aspartic acid and lysine), adenosine 5’ monophosphate monohydrate (AMP), adenosine, adenine, adenosine 5’ triphosphate (ATP), adenosine 2⍰, 3⍰-cyclic monophosphate (211, 311-cAMP), 1-ethyl-3-(3-dimethylaminopropyl) carbodiimide hydrochloride (EDC) and magnesium chloride (MgCl_2_) were procured from Sigma Aldrich. Standard glycine oligopeptide controls (dimer-hexamer), the HPLC buffer salts including sodium perchlorate and tris salts were also procured from Sigma Aldrich. Standard AMP oligonucleotide controls including dimer to tetramer of AMP i.e. AMP-AMP, AMP-AMP-AMP, AMP-AMP-AMP-AMP) were custom synthesised from Dharmacon™ siRNA solutions. The LCMS solvents, acetonitrile (can) and water, were purchased from J.T.Baker® while N,N-Dimethylformamide (DMF), triethyl amine (TEA) and glacial acetic were obtained from Finar. All reagents were used without further purification.

## Methods

### A. Dehydration-Rehydration (DH-RH) reaction setup

In a typical DH-RH reaction mixture of 300 uL volume, ATP/cAMP was added to a final concentration of 25mM. The amino acids (glycine/tryptophan/aspartic acid/lysine) were added to a final concentration of 25mM or 75mM (for only glycine). MgCl2 was added to a final concentration of 10mM in case of all reactions that comprised it. The pH was set to 7 (in all the reactions studied case of glycine and lysine) with 1M NaOH or 1.5N HCl, as need be. After setting the pH to 7, 0 time point sample (Cycle 0) was taken out. The rest of the reaction mix was transferred to 20 mL glass vials with their caps fitted with PTFE septa purchased from Chemglass. These were placed on a heating block set at elevated temperatures (90°C/ 130°C, depending on the experiment) to simulate early Earth geological conditions (Mungi et al., 2015). Predetermined time points were taken at 1hr (Cycle 1), 2hrs (Cycle 2), 5hrs (Cycle 5), 10hrs (Cycle 10) and 24hr (Cycle 24), after rehydration the dry sample with Milli Q water.

### B. Anion Exchange-HPLC analysis

HPLC analysis was carried out using an Agilent Technologies Infinity Series 1260 HPLC instrument. The time points from the DH-RH reaction were loaded onto the HPLC system after filtering them with 0.22µm Costar Spin-X filter centrifugation tubes to separate any solid impurities. Separation of molecules was achieved through the DNAPac PA 200 anion-exchange column from Thermo Scientific. This column separates the molecules based on the column matrix’s interaction with the negatively charged phosphate groups. Samples were analyzed using 20 mM Tris, pH 8 (Solvent A) and 400 mM NaClO4 in 20 mM Tris buffer, pH 8 (Solvent B) with a step gradient [0% Solvent B, 3 mins; 30% Solvent B, 10 mins; 100% Solvent B, 13 mins; 100% Solvent B, 16 mins; 0% Solvent B, 18 mins; 0% Solvent B, 21 mins] at a flow rate of 1.2 mL/min. Detection of analytes was carried out at 260 nm using a photo diode-array detector (DAD), using a highly sensitive 60 mm flow cell. A typical HPLC chromatogram revealed the presence of adenine, adenosine, cAMP monomer, linear AMP monomer, ATP, and oligomers. To semi-quantify the analytes, the area under the respective peaks was measured. It is worth noting that while the semi-quantified area could comprise of various species carrying the same charge (for example adenosine 5’-dihosphate and AMP-AMP molecules), it still provided insights into the distribution of species following a specific DH-RH cycle in the reaction.

### C. Liquid Chromatography Mass Spectrometry (LCMS)

The mass analysis of the filtered samples was achieved utilizing a Sciex X500R QTOF mass spectrometer (MS), coupled with an Exion-LC series UHPLC system from Sciex. The scanning approach employed was information-dependent acquisition (IDA). The initial reaction mixture was subjected to separation on a Zorbax C8 column [Dimensions: 4.6 × 150 mm, particle size: 3.6 μm] supplied by Agilent technologies. A gradient of mixture of LCMS grade water containing 0.1% formic acid and acetonitrile containing 0.1% formic acid was employed for the separation of the oligomers. Each run time was of 25 mins with a flow rate of 0.5 ml/mins. For TOF-MS analysis, Electron spray ionization (ESI) was used for all mass acquisitions, using the following specific instrument parameters; ion source gas 1 & 2 was kept at 40 and 50 psi respectively, curtain gas 30 L/min, CAD gas 7 psi, ion spray voltage set at 5500 V (in positive mode) and an operating temperature of 500°C. The acquisition for Time-of-Flight Mass Spectrometry (TOF-MS) was conducted at an 80 V declustering potential, with a spread of 20 V, and a collision energy of 10 V. An internal standard (alanine, 20mg/ml) was injected with 150 nmoles of samples to obtain the yields of resultant products, which were then summed up to get the overall oligomer yield. All quantitation experiments were performed in at least 6 replicates. Analysis of the acquired data was done using Sciex OS software, from University of Florida. The confirmation of a specific species or molecule’s presence involved verifying the precursor mass within a 5-ppm error range. For TOF MS–MS analysis, a collision energy of 50 V with a spread of 20 V was employed. Since mass acquisition was conducted in the positive mode, the observed masses corresponded to the mass of the H+ adduct of the parent molecule. The confirmation of a specific molecule was identified using its signature fragmentation pattern mass within a 20-ppm error range.

For low intensity peaks of compounds, for e.g., AMP tetramer (AMP) 4, Agilent MS Q-TOF Dual AJS ESI (component model, G6545B) was utilized for TOF MS-MS spectra. 10 µl (5nmoles) of filtered samples was subjected to separation on a Phenomenex Luna C18 Column [Dimension: 250 * 4.6 mm, particle size: 5 µm and pore size: 100 Å] with column oven temperature set at 40°C. Mass spectra were acquired in the positive ion polarity using solvents containing gradient mixture of LCMS grade water containing 0.1% formic acid and methanol containing 0.1% formic acid (again, gradient details in brief along with run time and flow rate). For the separation of the oligomers with specific instrument parameters [Gas Temp 320°C, Gas Flow 11 L/min, Nebulizer 45 psig, Sheath Gas Temperature 320°C and Sheath Gas Flow 11] and scanning parameters [VCap 3000, Nozzle Voltage 1000 V, Fragmentor voltage 80 and collision energy formula set with slope 2 and offset of 5] was standardized. For analysis, Agilent Mass Hunter Qualitative analysis software [Version 10.0] was used and molecule specific spectra were acquired using a Find by Formula approach.

## Acknowledgements

This research was supported by grants from the Science and Engineering Research Board (SERB), Department of Science and Technology, Government of India [CRG/2021/001851] and IISER Pune. The authors acknowledge DST-FIST grant SR/FST/LSII-043/2016 that enabled the establishment of the LC-MS facility at IISER Pune. RR thanks the Department of Biotechnology, Government of India for fellowship support. The authors would like to especially thank the lab members for their critical feedback.

## Authors contribution

RR and SR designed the experiments while RR performed the experiments. RR, AS and SR analysed the mass spectrometry and HPLC data. AS helped with the ChemDraw structures. RR and SR wrote the manuscript with relevant inputs from AS.

## Corresponding authors

Correspondence to Dr. Sudha Rajamani

## Ethics declarations

### Competing interests

The authors declare no competing interests

## Notes

### Competing Interest Statement

The authors have declared no competing interest.

## References

1. Fahrenbach, A. C., Giurgiu, C., Tam, C. P., Li, L., Hongo, Y., Aono, M., & Szostak, J. W. Common and Potentially Prebiotic Origin for Precursors of Nucleotide Synthesis and Activation. Journal of the American Chemical Society, 139, 8780–8783 (2017).

2. Mariani, A., Russell, D. A., Javelle, T., & Sutherland, J. D. A Light-Releasable Potentially Prebiotic Nucleotide Activating Agent. Journal of the American Chemical Society, 140, 8657–8661 (2018).

3. Stolar, T., Grubešić, S., Cindro, N., Meštrović, E., Užarević, K., & Hernández, J. G. Mechanochemical Prebiotic Peptide Bond Formation. Angewandte Chemie International Edition, 60, 12727–12731(2021).

4. Leman, L., Orgel, L., & Ghadiri, M. R. (2004). Carbonyl Sulfide-Mediated Prebiotic Formation of Peptides. Science, 306, 283–286.

5. Rode, B. M., & Schwendinger, M. G. Copper-catalyzed amino acid condensation in water — A simple possible way of prebiotic peptide formation. Origins of Life and Evolution of the Biosphere, 20, 401–410 (1990).

6. Liu, Z., Wu, L.-F., Xu, J., Bonfio, C., Russell, D. A., & Sutherland, J. D. Harnessing chemical energy for the activation and joining of prebiotic building blocks. Nature Chemistry, 12, 1023–1028 (2020).

7. Koshland, D. E. Kinetics of Peptide Bond Formation. Journal of the American Chemical Society, 73, 4103–4108 (1951).

8. Peller, L. The co-association of nucleosides and the equilibrium copolymerization of nucleotides. Base stacking interactions and the thermodynamics of phosphodiester bond formation. The Journal of Physical Chemistry, 80, 2462–2467 (1976).

9. Rodriguez-Garcia, M., Surman, A. J., Cooper, G. J. T., Suárez-Marina, I., Hosni, Z., Lee, M. P., & Cronin, L. Formation of oligopeptides in high yield under simple programmable conditions. Nature Communications, 6, 8385 (2015).

10. Dass, A. V., Wunnava, S., Langlais, J., von der Esch, B., Krusche, M., Ufer, L., Chrisam, N., Dubini, R. C. A., Gartner, F., Angerpointner, S., Dirscherl, C. F., Rovó, P., Mast, C. B., Šponer, J. E., Ochsenfeld, C., Frey, E., & Braun, D. RNA Oligomerisation without Added Catalyst from 2′,3′-Cyclic Nucleotides by Drying at Air-Water Interfaces. ChemSystemsChem, 5 (2023).

11. Müller, F., Escobar, L., Xu, F., Węgrzyn, E., Nainytė, M., Amatov, T., Chan, C. Y., Pichler, A., & Carell, T. A prebiotically plausible scenario of an RNA–peptide world. Nature. 605, 279–284 (2022).

12. Bapat, N. v., & Rajamani, S. (2015). Effect of Co-solutes on Template-Directed Nonenzymatic Replication of Nucleic Acids. Journal of Molecular Evolution, 81, 72– 80 (2015).

13. Patki, G. M., & Rajamani, S. Nonenzymatic RNA replication in a mixture of ‘spent’ nucleotides. FEBS Letters, 597, 3125–3134 (2023).

14. DasGupta, S., Zhang, S., & Szostak, J. W. Molecular Crowding Facilitates Ribozyme-Catalyzed RNA Assembly. ACS Central Science, 9, 1670–1678 (2023).

15. Joshi, M. P., Sawant, A. A., & Rajamani, S. Spontaneous emergence of membrane-forming protoamphiphiles from a lipid–amino acid mixture under wet–dry cycles. Chemical Science, 12, 2970–2978 (2021).

16. Sarkar, S., Dagar, S., Verma, A., & Rajamani, S. (2020). Compositional heterogeneity confers selective advantage to model protocellular membranes during the origins of cellular life. Scientific Reports, 10, 4483 (2020).

17. Patel, B. H., Percivalle, C., Ritson, D. J., Duffy, C. D., & Sutherland, J. D. Common origins of RNA, protein and lipid precursors in a cyanosulfidic protometabolism. Nature Chemistry, 7, 301–307 (2015).

18. Chu, X.-Y., & Zhang, H.-Y. Prebiotic Synthesis of ATP: A Terrestrial Volcanism-Dependent Pathway. Life, 13, 731 (2023).

19. Biscans, A. Exploring the Emergence of RNA Nucleosides and Nucleotides on the Early Earth. Life, 8, 57 (2018).

20. Kim, H.-J., & Benner, S. A. Abiotic Synthesis of Nucleoside 5′-Triphosphates with Nickel Borate and Cyclic Trimetaphosphate (CTMP). Astrobiology, 21, 298–306 (2021).

21. Leslie E. O., Prebiotic Chemistry and the Origin of the RNA World. Critical Reviews in Biochemistry and Molecular Biology, 39, 99–123 (2004).

22. Mohamady, S., & Taylor, S. D. One flask synthesis of 2′,3′-cyclic nucleoside monophosphates from unprotected nucleosides using activated cyclic trimetaphosphate. Tetrahedron Letters, 57, 5457–5459 (2016).

23. Yamagata, Y., Inoue, H., & Inomata, K. Specific effect of magnesium ion on 2′, 3′-cyclic amp synthesis from adenosine and trimeta phosphate in aqueous solution. Origins of Life and Evolution of the Biosphere, 25, 47–52 (1995).

24. Jiang, L., Dziedzic, P., Spacil, Z., Zhao, G.-L., Nilsson, L., Ilag, L. L., & Córdova, A. Abiotic synthesis of amino acids and self-crystallization under prebiotic conditions. Scientific Reports, 4, 6769 (2014).

25. Ring, D., Wolman, Y., Friedmann, N., & Miller, S. L. Prebiotic Synthesis of Hydrophobic and Protein Amino Acids. Proceedings of the National Academy of Sciences, 69, 765–768 (1972).

26. Breslow, R. Formation of L Amino Acids and D Sugars, and Amplification of their Enantioexcesses in Aqueous Solutions, Under Simulated Prebiotic Conditions. Israel Journal of Chemistry, 51, 990–996 (2011).

27. Pinna, S., Kunz, C., Halpern, A., Harrison, S. A., Jordan, S. F., Ward, J., Werner, F., & Lane, N. A prebiotic basis for ATP as the Universal Energy Currency. PLOS Biology, 20 (2022).

28. Jiménez, E. I., Gibard, C., & Krishnamurthy, R. Prebiotic Phosphorylation and Concomitant Oligomerization of Deoxynucleosides to form DNA. Angewandte Chemie International Edition, 60, 10775–10783 (2021).

29. Kim, H.-J., & Benner, S. A. Abiotic Synthesis of Nucleoside 5′-Triphosphates with Nickel Borate and Cyclic Trimetaphosphate (CTMP). Astrobiology, 21(3), 298–306 (2021).

30. Dagar, S., Sarkar, S., & Rajamani, S. Geochemical influences on nonenzymatic oligomerization of prebiotically relevant cyclic nucleotides. RNA, 26, 756–769 (2020).

31. Sakata, K., Yabuta, H., & Kondo, T. Effects of metal ions and pH on the formation and decomposition rates of di- and tri-peptides in aqueous solution. Geochemical Journal, 48, 219–230 (2014).

32. Mullins, D. W., & Lacey, J. C. Studies of the chemical basis of the origin of protein synthesis: Initiation and direction of peptide growth. Journal of Molecular Evolution, 15, 339–345 (1980).

33. Wu, L. F., Su, M., Liu, Z., Bjork, S. J., & Sutherland, J. D. Interstrand Aminoacyl Transfer in a tRNA Acceptor Stem-Overhang Mimic. Journal of the American Chemical Society, 143, 11836–11842 (2021).

34. Ni, F., Fu, C., Gao, X., Liu, Y., Xu, P., Liu, L., Lv, Y., Fu, S., Sun, Y., Han, D., Li, Y.-M., & Zhao, Y. N-phosphoryl amino acid models for P-N bonds in prebiotic chemical evolution. Science China Chemistry. 58, 374–382 (2015).

35. Pascal, R., & Boiteau, L. Energy flows, metabolism and translation. Philosophical Transactions of the Royal Society B: Biological Sciences, 366, 2949–2958 (2011).

36. Jahansouz, H., Jiang, Z., Himes, R. H., Mertes, M. P., & Mertes, K. B. Formate activation in neutral aqueous solution mediated by a polyammonium macrocycle. Journal of the American Chemical Society, 111, 1409–1413 (1989).

37. Misra, V. K., & Draper, D. E. On the role of magnesium ions in RNA stability. Biopolymers: Original Research on Biomolecules, 48, 113–135 (1998).

38. Yamagami, R., Bingaman, J. L., Frankel, E. A., & Bevilacqua, P. C. Cellular conditions of weakly chelated magnesium ions strongly promote RNA stability and catalysis. Nature communications, 9, 2149 (2018).

39. Sissi, C., & Palumbo, M. Effects of magnesium and related divalent metal ions in topoisomerase structure and function. Nucleic Acids Research, 37(3), 702–711(2009).

40. Carvalho, A. T. P., Fernandes, P. A., & Ramos, M. J. The Catalytic Mechanism of RNA Polymerase II. Journal of Chemical Theory and Computation, 7, 1177–1188 (2011).

41. Roberts, S. J., Liu, Z., & Sutherland, J. D. Potentially Prebiotic Synthesis of Aminoacyl-RNA via a Bridging Phosphoramidate-Ester Intermediate. Journal of the American Chemical Society, 144, 4254–4259 (2022).

42. Su, M., Schmitt, C., Liu, Z., Roberts, S. J., Liu, K. C., Röder, K., Jäschke, A., Wales, D. J., & Sutherland, J. D. Triplet-Encoded Prebiotic RNA Aminoacylation. Journal of the American Chemical Society, 145, 15971–15980 (2023).

43. Singer, J. N., Müller, F. M., Węgrzyn, E., Hölzl, C., Hurmiz, H., Liu, C., Escobar, L., & Carell, T. Loading of Amino Acids onto RNA in a Putative RNA-Peptide World. Angewandte Chemie International Edition, 62, e202302360 (2023).

44. Radakovic, A., Lewicka, A., Todisco, M., Aitken, H. R. M., Weiss, Z., Kim, S., Bannan, A., Piccirilli, J. A., & Szostak, J. W. A potential role for RNA aminoacylation prior to its role in peptide synthesis. Proceedings of the National Academy of Sciences of the United States of America, 121, e2410206121 (2024).

